# A homozygous human *WNT11* loss-of-function variant associated with laterality, heart and renal defects

**DOI:** 10.1101/2024.11.14.623711

**Authors:** Henrike Berns, Maximilian Haas, Zeineb Bakey, Magdalena Maria Brislinger-Engelhardt, Miriam Schmidts, Peter Walentek

## Abstract

Wnt signaling plays important roles during vertebrate development, including left-right axis specification as well as heart and kidney organogenesis. We identified a homozygous human *WNT11* variant in an infant with *Situs inversus totalis*, complex heart defects and renal hypodysplasia, and we used *Xenopus* embryos to functionally characterize this variant. *WNT11^c.814delG^* encodes a loss-of-function protein with reduced stability that lost signaling activity *in vivo*. This is remarkable, because the variant encodes a truncated ligand with nearly identical length and predicted structure to dominant-negative Wnts. Furthermore, we demonstrate that alteration of the truncated C-terminal end can restore stability and dominant-negative signaling activity. Our study also suggests similar functions for WNT11 in human development as described in model organisms. Therefore, biallelic WNT11 dysfunction should be considered as novel genetic cause in syndromal human phenotypes presenting with congenital heart defects and renal hypoplasia, with or without laterality defects. The work presented here enhances our understanding of human development and structure-function relationships in Wnt ligands.

## Introduction

Bilaterians have an overall bilateral symmetric body plan, but several internal organs display asymmetries along the left-right body axis (Fliegauf et al., 2007; Namigai et al., 2014). The most common organ arrangement is called *situs solitus*, where in humans, the lung consists of two lobes on the left and three on the right side, the liver is positioned on the right side and the heart is tilted towards the left side (Fliegauf et al., 2007). Laterality defects, mis-arrangements of left-right asymmetric organs, occur in about 1 in 10,000 human live births (Afzelius, 1999; Lin et al., 2014; Torgersen, 1950). These include *situs inversus totalis* (SIT), heterotaxia and isomerisms (Fliegauf et al., 2007). While SIT is a deviation from the norm without direct health consequence, heterotaxia and isomerisms can result in misplacements, duplications or absence of individual organs, congenital heart defects (CHDs) as well as structural defects of the great vessels (Fakhro et al., 2011; Lin et al., 2014).

Organ asymmetries are initiated during development by the left-right organizer (LRO), which breaks bilateral symmetry leading to asymmetric gene expression guiding further organ morphogenesis (Blum et al., 2014; Hamada, 2020). The LRO is a monociliated epithelium and its transient appearance, structure and function are remarkably conserved across most vertebrate species (Blum et al., 2009b). In the frog *Xenopus laevis*, the LRO is called gastrocoel roof plate (GRP) (Schweickert et al., 2007; Shook et al., 2004). Central cells of the GRP project motile cilia, which are asymmetrically positioned at the posterior poles of cells and beat in a clockwise fashion to generate an extracellular leftward fluid flow (Walentek et al., 2012). Cells in the lateral margins of the GRP harbor non-motile sensory cilia required to sense fluid flow (Gopalakrishnan et al., 2023; Schweickert et al., 2007). Lateral GRP cells also express the signaling ligand Nodal and its inhibitor Dand5 (also called Coco in *Xenopus*) (Schweickert et al., 2010). Leftward flow reduces *coco* expression, leading to release of Nodal repression exclusively on the left side. Nodal then induces an asymmetric gene expression cascade in the left lateral plate mesoderm by activating its own expression as well as expression of its feedback inhibitor *lefty* and the transcription factor *pitx2c* (Blum et al., 2014; Schweickert et al., 2001; Schweickert et al., 2010). Pitx2c remains active on the left side of the embryo during subsequent asymmetric organ development.

*Xenopus* is particularly suited to study early development and laterality defects, because embryos develop externally, are large in size, gastrulation stages are reached within a few hours, and tissues such as the left or right side of the GRP can be selectively targeted by micro-injection and manipulations (Blum et al., 2009a; Walentek et al., 2012).

In humans, laterality defects can occur as a consequence of defective fluid due to missing or malformed LRO cilia, for which a vast number of responsible genes have been identified (Antony et al., 2022; Fliegauf et al., 2007; Raidt et al., 2023; Reiter and Leroux, 2017). Furthermore, defects in genes related to cell signaling pathways have also been found to result in laterality defects (Minegishi et al., 2023; Mohapatra et al., 2009; Sempou and Khokha, 2019). Nevertheless, causative genes for laterality defects can only be identified in less than half of the affected human individuals (Antony et al., 2022).

Wnt is a signaling pathway with multiple branches affecting gene expression (e.g. canonical Wnt/β-catenin pathway), cell polarity (e.g. Wnt/planar cell polarity (PCP) pathway) and morphogenesis (e.g. Wnt/calcium pathway) (Minegishi et al., 2023; Walentek et al., 2012; Walentek et al., 2018; Walentek et al., 2013). In *Xenopus* development, Wnt/β-catenin is required for dorso-ventral axis induction (a prerequisite for left-right axis development) and for generation of motile GRP cilia by activating the transcription factor *foxj1* (Walentek et al., 2012). Wnt/PCP signaling is required for posterior positioning of GRP cilia required to generate an asymmetric leftward fluid flow (Walentek et al., 2012). Other Wnt pathway branches affect gastrulation movements and normal GRP morphogenesis, which has also been shown to affect the development of motile vs. non-motile GRP cilia and lateral GRP gene expression (Chien et al., 2018; Kühl et al., 2000; Tada and Smith, 2000; Walentek et al., 2013).

Particularly, *Xenopus* Wnt11b has been identified as a regulator of gastrulation movements (Tada and Smith, 2000), GRP morphology, Wnt/PCP and left-right axis formation (Walentek et al., 2013). Wnt11b loss-of-function in the GRP also reduced expression of *nodal1* and *coco*, indicating that Wnt11b affects left-right symmetry breakage primarily by interfering with GRP morphogenesis and the signaling cascade downstream of the cilia-dependent symmetry breakage (Walentek et al., 2013). The association of Wnt11b with laterality has recently been confirmed in *Xenopus wnt11b^-/-^* knockout animals (Houston et al., 2022). Notably, Wnt11 is also expressed in the LROs of other species, such as the Henseńs node in chicken and the node of mice, suggesting a role of Wnt11 in the determination of left-right asymmetry across vertebrates (Chapman et al., 2004; Kispert et al., 1996).

Besides regulating left-right axis development, Wnt11 also functions in other organ systems. Overexpression of a dominant-negative (dn)Wnt11b induced kidney developmental defects in *Xenopus* (Dichmann et al., 2015), and *Wnt11^-/-^* mice develop hypoplastic kidneys and CHDs with complete penetrance (Majumdar et al., 2003; Zhou et al., 2007). Thus, Wnt11b has been related to laterality defects in *Xenopus*, and knock-out of *Wnt11* has been identified as a cause for heart and kidney defects in mice.

Here, we have identified a homozygous human *WNT11* variant (c.814delG, p.Glu272Asn*13) in an infant with SIT, Tetralogy of Fallot (TOF) and severe bilateral renal hypodysplasia. A biallelic *WNT11* variant has to our knowledge not been identified in humans before, and WNT11 dysfunction has not previously been identified as a cause of these human developmental defects. Based on this discovery, we used *Xenopus* embryos as model system to functionally characterize this novel *WNT11* variant. We found that *WNT11^c.814delG^* encodes a truncated ligand with reduced stability, which lost its ability to activate Wnt signaling as well as to regulate morphogenesis. Thereby, the resulting protein functionally differs from dominant-negative acting truncations of similar length described in the literature, despite only minor differences in the amino acid sequence. Together, we have identified a new human *WNT11* variant associated with a syndromale developmental phenotype, functionally characterized the altered WNT11 protein, and discovered how differences in the WNT11 protein can dramatically alter its functions.

## Results

### Identification of a homozygous *WNT11* variant in an individual with laterality, cardiac and renal defects

We performed exome sequencing including Copy Number Variant (CNV) analysis in a first-born infant of consanguineous south-east Asian descent with SIT including dextrocardia, in combination with a complex congenital heart defect (TOF: ventricular septal defect, overriding aorta, pulmonary stenosis and right ventricular hypertrophy) and severe bilateral renal hypodysplasia, with end-stage renal disease (ESRD) in infancy. This revealed a homozygous *WNT11* frameshift variant, c.814delG, p.Glu272Asn*13 (**Fig. 1A; S1A**). The variant was absent from gnomAD (gnomad.broadinstitute.org/) with only a missense variant at the same amino acid position 272, Glu272Lys, reported in 1 of 1,614,052 alleles. In total, as of October 2024, there are 38 different stop or frameshift variants in *WNT11* reported in gnomAD. All of these are heterozygous and the majority represent private alleles, all with a very low allele frequency (< 0.0001 minor allele frequency), suggesting that WNT11 has non-redundant functions and that biallelic loss-of-function variants will result in a human phenotype. In the affected individual, no additional putative disease-causing variants or CNVs were detected. This indicated the *WNT11^c.814delG^* variant as likely cause in our individual and suggested WNT11 dysfunction as novel cause for congenital defects in humans.

**Figure 1:**
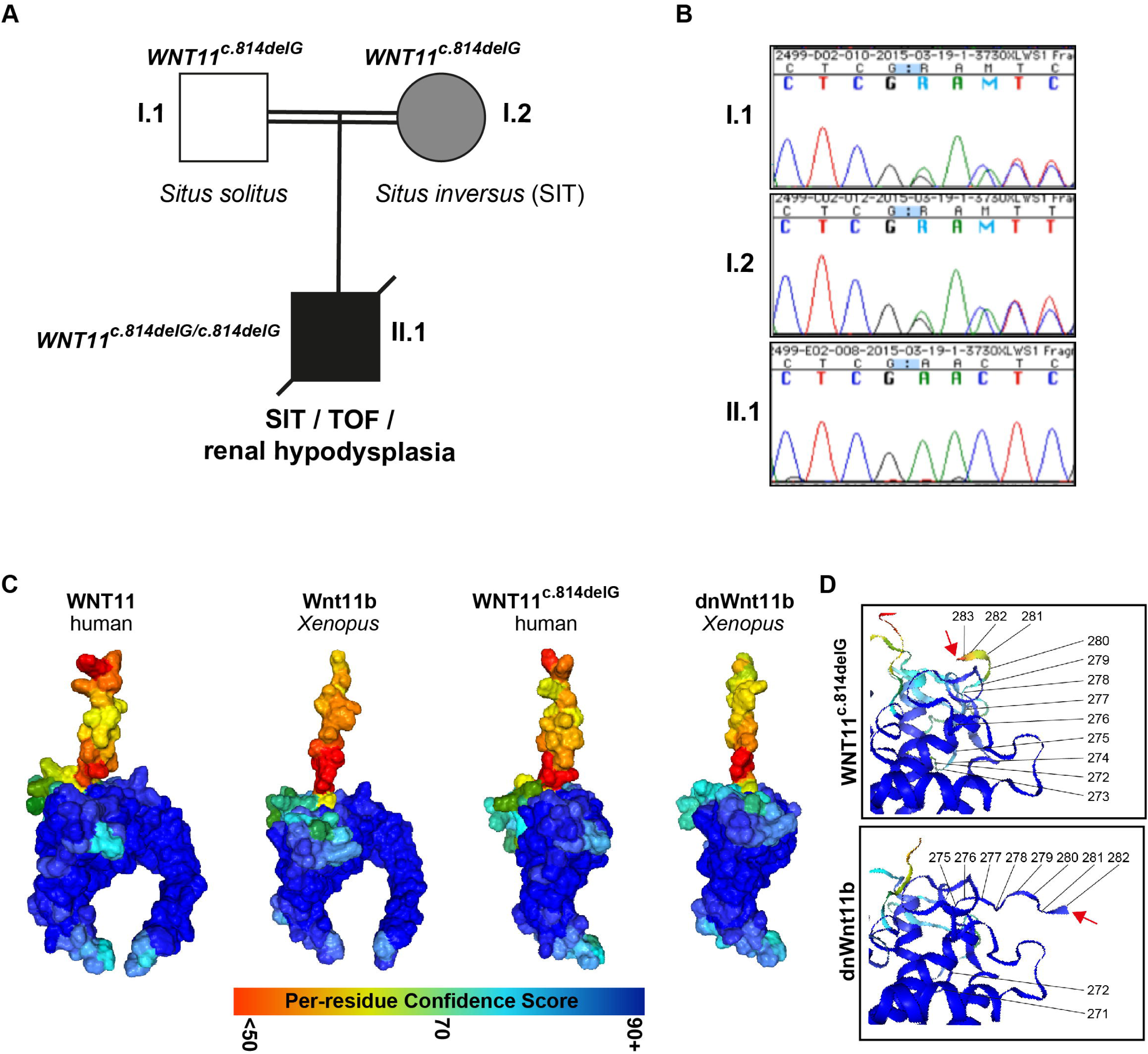
A *WNT11* variant associated with laterality, heart and kidney developmental defects. **(A)** Pedigree indicating clinical features and segregation of the identified *WNT11^c.814delG^* variant amongst the three family members. **(B)** Chromatogram traces of Sanger sequencing results regarding the *WNT11* variant segregation within the family, showing a homozygous deletion of c.814G in the affected child while both parents are heterozygous. In **(A,B)**, I.1 = father; I.2 = mother; II.1 = son. **(C,D)** Alphafold 2.1.2 protein structure predictions of human (WNT11) and *Xenopus* (Wnt11b) Wnt ligands, and comparison the WNT11^c.814delG^ variant to dominant-negative (dn)Wnt11b. **(D)** Magnification of C-termini of truncated ligands with annotation of the last amino acids. Red arrows indicate C-terminal end.

To investigate if *WNT11* variants could be found in additional patients with overlapping phenotypes, we performed sanger sequencing of *WNT11* coding and splice site regions (positions +/-20) of 39 cases with non-cystic renal dysplasia, 13 cases of severe renal hypoplasia, and one case of renal hypoplasia and heart defect. This did not reveal any putative causative *WNT11* variants. Furthermore, we did not identify any additional likely disease-causing *WNT11* variants using exome sequencing in a cohort of 37 cases with laterality defects (Antony et al., 2022) nor in 3 additional cases with renal hypoplasia. Likewise, we did not identify any additional cases with biallelic *WNT11* variants through our national and international collaborators or through a genematcher search (https://genematcher.org/).

Interestingly, the mother of the affected child also presented with SIT while the father was not affected. Segregation analysis within the family revealed that both parents were heterozygous for the *WNT11^c.814delG^* frameshift allele (**Fig. 1A,B**). To exclude an additional underlying monogenic cause for the laterality defect in the mother, we performed exome sequencing also for her. However, this did not reveal any additional plausible genetic causes.

We investigated the predicted protein structures of wt WNT11 and WNT11^c.814delG^ using Alphafold v2.1.2 (Jumper et al., 2021), which revealed the stereotypical hand-like structure of the wt Wnt ligand (Willert and Nusse, 2012), while the ligand encoded by *WNT11^c.814delG^* was lacking the C-terminal “index finger” in the protein structure (**Fig. 1C**). Interestingly, C-terminally truncated Wnt ligands were previously shown to exhibit dominant-negative (dn) effects on Wnt signalling *in vivo*, including an engineered *Xenopus laevis* dnWnt11b construct that affected gastrulation movements, morphogenesis and laterality in embryos (Hoppler et al., 1996; Tada and Smith, 2000; Walentek et al., 2013). Therefore, we compared wt human WNT11 and *Xenopus* Wnt11b structures, which confirmed the expected high degree of similarity (**Fig. 1C**). In contrast, comparison of dnWnt11b and WNT11^c.814delG^ predicted structures suggested that changes in the amino acid sequence due to the frameshift in WNT11^c.814delG^ altered the structural prediction for the last 12 amino acids with unknown consequences for protein function (**Fig. 1C,D**).

In summary, these results revealed a novel homozygous human *WNT11* variant associated with complex developmental defects and predicted structural alterations of the encoded Wnt ligand.

### Functional testing of *WNT11^c.814delG^* effects on embryonic development in *Xenopus*

Two Wnt11 paralogs exist in *Xenopus*: Wnt11b is maternally deposited, expressed in the superficial layer of the gastrula organizer and the blastopore as well as the developing somites; Wnt11 (also called Wnt11r) is expressed in the developing nervous system and the cement gland (Dichmann et al., 2015; Garriock et al., 2005; Matthews et al., 2008; Tada and Smith, 2000; Walentek et al., 2013). Wnt11 functions in development have been extensively studied in *Xenopus* embryos, and Wnt11b has been shown to regulate gastrulation movements, morphogenesis as well as left-right axis specification after both gain- and loss-of-function (Walentek et al., 2013). We therefore chose to test how human WNT11 and WNT11^c.814delG^ affect early *Xenopus* development in comparison to *Xenopus* Wnt11b and dnWnt11b *in vivo*.

For that, we injected mRNAs encoding Wnt11b, dnWnt11b, WNT11 and WNT11^c.814delG^ (**Fig. S1A**) targeting the prospective dorsal mesoderm, and analyzed the impact on morphogenesis in gastrula and neurula (st. 13 and 20) embryos. As previously described, overexpression of *Xenopus* full-length Wnt11b and dnWnt11b impaired gastrulation movements leading to high frequencies of blastopore and neural tube closure defects (> 70%) (**Fig. 2A,B**) (Walentek et al., 2013). Overexpression of full-length WNT11 induced morphological defects at similar frequencies to Wnt11b, confirming a high degree of functional homology between Wnt11b and WNT11 (**Fig. 2A,B**). In contrast, overexpression of *WNT11^c.814delG^* induced morphological defects at much lower frequencies (< 40%), indicating functional differences to all other constructs, including dnWnt11b (> 70%) (**Fig. 2A,B**). Mortality rates in embryos were similar across manipulations, excluding the possibility that fewer morphological defects were recovered after *WNT11^c.814delG^* overexpression due to early embryonic lethality (**Fig. S1B**).

**Figure 2:**
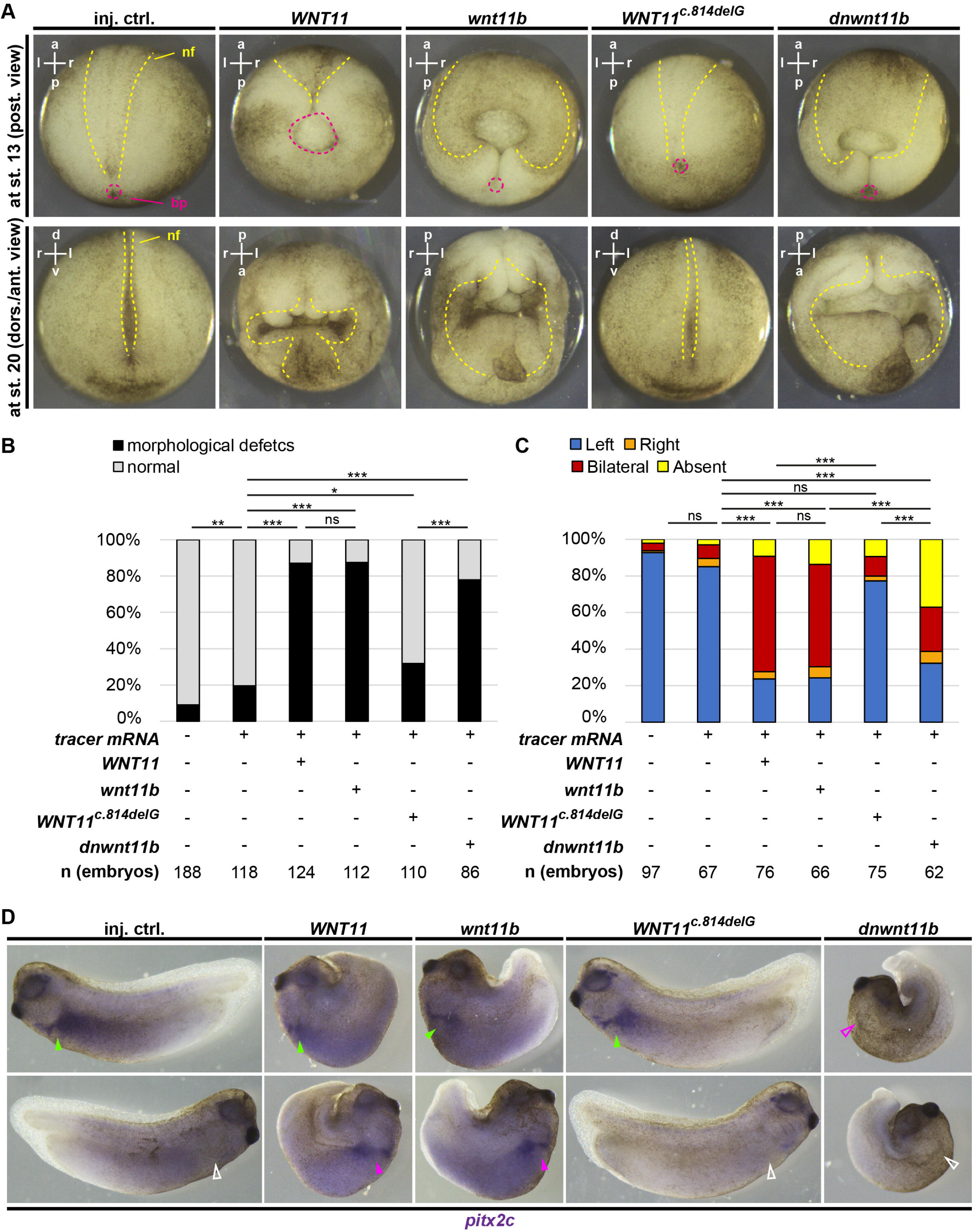
Functional testing of the WNT11^c.814delG^ variant in *Xenopus* embryos. **(A-D)** Overexpression of mRNAs encoding WNT11, Wnt11b, WNT11^c.814delG^ and dnWnt11b, and analysis of effects on **(A,B)** morphogenesis at st. 13 (posterior view) and 20 (dorso-anterior view), **(C,D)** and left-right axis patterning (lateral views) by WMISH at st. 30 - 32 against *pitx2c*. **(A,D)** Representative images. In **(A)**, neural folds (nf) and blastopores (bp) are indicated by yellow and magenta dashed lines, respectively. In **(D)**, filled green arrowheads indicate normal left-sided expression, open white arrowheads indicate normal right-sided lack of expression. Filled magenta arrowheads indicate ectopic right-sided expression, open magenta arrowheads indicate loss of left-sided expression. **(B,C)**, Quantification of results. n = number of embryos analyzed per condition. *χ*^2^-test, p > 0.05 = ns; p < 0.05 = *; p < 0.01 = **; p < 0.001 = ***.

Next, we tested whether WNT11 and WNT11^c.814delG^ overexpression can cause laterality defects in *Xenopus*. For that, *Wnt11b*, *dnWnt11b*, *WNT11* and *WNT11^c.814delG^* mRNAs were targeted to the dorsal mesoderm and LRO (Schweickert et al., 2012; Walentek et al., 2012), and left-right axis specification was analyzed using asymmetric *pitx2c* gene expression at st. 30 - 32 by whole mount *in situ* hybridization (WMISH). In control embryos, *pitx2c* expression was found to be predominantly (> 80%) expressed exclusively on the left side of the embryo (**Fig. 2C,D**). Gain of wt Wnt11b or WNT11 induced ectopic right-sided *pitx2c* leading to bilateral expression in the majority of embryos (> 50% bilateral), while dnWnt11b caused a loss of *pitx2c* expression as most frequent phenotype (> 30% absent) (**Fig. 2C,D**), in line with published results (Walentek et al., 2013). Importantly, overexpression of WNT11^c.814delG^not only led to embryos developing no visible morphological defects, but also no left-right axis defects were detected in the majority of manipulated embryos (> 70% left-sided expression) (**Fig. 2C,D**).

Taken together, *in vivo* functional tests strongly suggest that WNT11^c.814delG^ encodes a hypomorph or loss-of-function ligand that lost its ability to activate Wnt signaling. Hence, overexpression of WNT11^c.814delG^ did not affect embryonic morphogenesis or left-right axis specification.

### Amino acid changes in *WNT11^c.814delG^* reduce protein stability and signaling properties

Structural predictions suggested differences between dnWnt11b and WNT11^c.814delG^, and *in vivo* assays indicated that *WNT11^c.814delG^* encodes a ligand which lost signaling ability. This was remarkable, because various similar C-terminal truncations of Wnt ligands were shown to generate dominant-negative acting signaling molecules (Dichmann et al., 2015; Hoppler et al., 1996; Tada and Smith, 2000). We therefore wondered whether the differences in length or differences in the amino acid sequence were the cause of the observed functional differences between dnWnt11b and WNT11^c.814delG^.

To investigate the molecular basis of functional differences between truncated Wnt ligands, we generated *WNT11* constructs (a.) lacking the frame-shifted portion (*WNT11^Δ814-1062^*) or (b.) with a corrected sequence, but same extent of truncation (*WNT11^Δ850-1062^*) as WNT11^c.814delG^ (**Fig. 3A, S1C**). We also generated (c.) a shortened *Xenopus* dnWnt11b construct (*Wnt11b^Δ811-1059^*) analogous to *WNT11^Δ814-1062^* (**Fig. 3A, S1C**) for comparison. mRNAs encoding the newly generated constructs were targeted to the dorsal mesoderm and LRO, and effects on asymmetric *pitx2c* expression as well as morphology were compared to effects induced by WNT11^c.814delG^and dnWnt11b. Like in previous experiments, WNT11^c.814delG^overexpression did not affect asymmetric *pitx2c* expression or embryonic morphogenesis as compared to controls (**Fig. 3B-E**). In striking contrast, removal of frameshift amino acids (*WNT11^Δ814-1062^*) generated results comparable to dnWnt11b (> 50% absent *pitx2c* expression and > 70% of embryos with morphological defects) (**Fig. 3B-E**). Correction of the frameshift amino acids (*WNT11^Δ850-1062^*) or shortening of the dnWnt11b ligand (*Wnt11b^Δ811-1059^*) both caused left-right and morphological defects, however at lower rates than dnWnt11b or WNT11^Δ814-1062^ (**Fig. 3B-E**). Thus, these results revealed striking functional differences in Wnt11-type ligands that differed only in few amino acids at their truncated C-termini.

**Figure 3:**
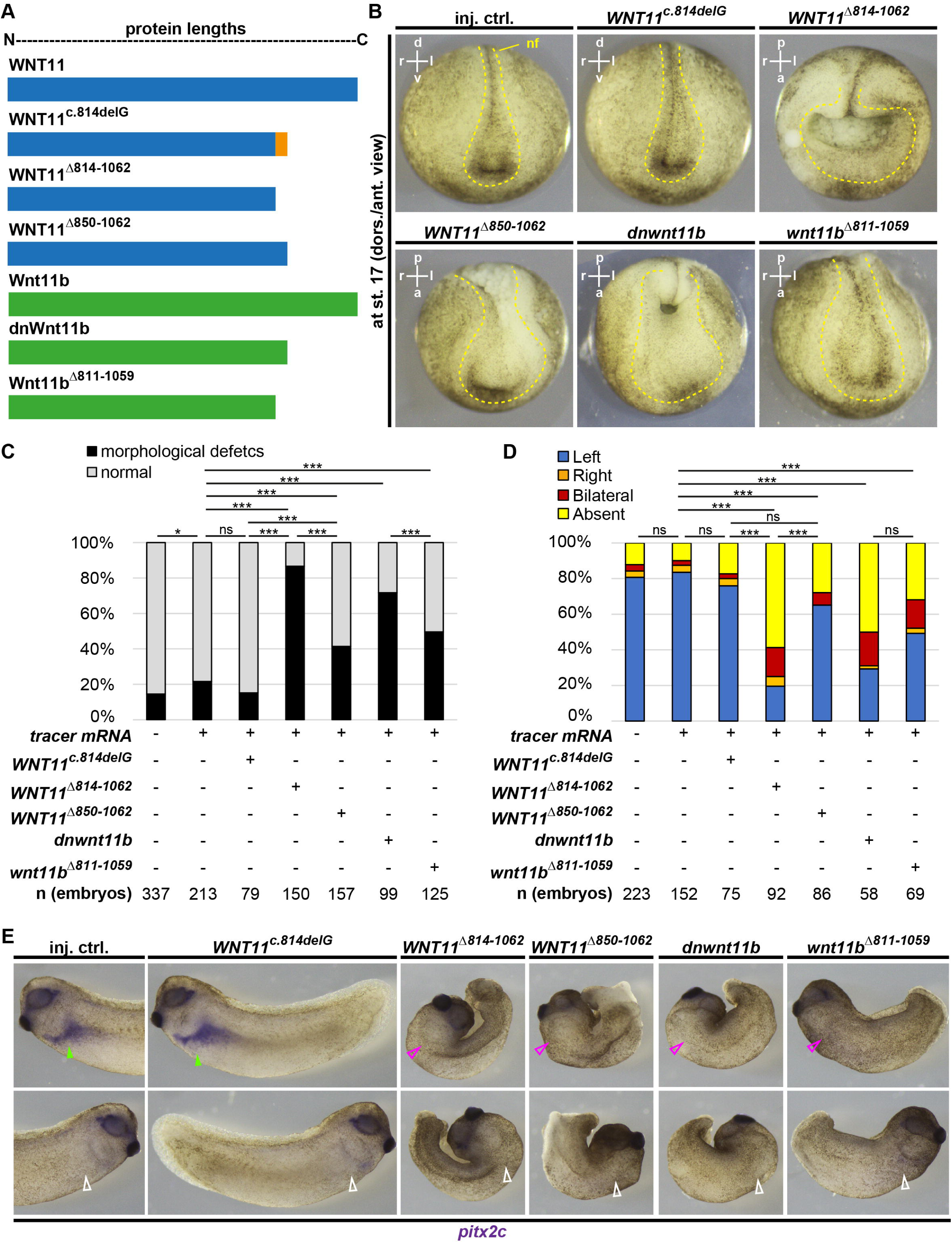
Changes in C-terminal amino acids alters WNT11^c.814delG^ function. **(A)** Schematic representation of constructs and their lengths. Human WNT11 constructs are shown in blue, *Xenopus* Wnt11b constructs are shown in green. Orange indicates changes in the sequence. **(B-E)** Overexpression of mRNAs encoding WNT11, WNT11^c.814delG^, WNT11^D814-1062^, WNT11^D850-1062^, dnWnt11b and Wnt11b^D811-1059^, and analysis of effects on **(B,C)** morphogenesis at st. 17 (dorso-anterior view), and **(D,E)** left-right axis patterning (lateral views) by WMISH at st. 30 - 32 against *pitx2c*. **(B,E)** Representative images. In **(B)**, neural folds (nf) are indicated by yellow dashed lines. In **(E)**, filled green arrowheads indicate normal left-sided expression, open white arrowheads indicate normal right-sided lack of expression. Filled magenta arrowheads indicate ectopic right-sided expression, open magenta arrowheads indicate loss of left-sided expression. **(C,D)** Quantification of results. n = number of embryos analyzed per condition. *χ*^2^-test, p > 0.05 = ns; p < 0.05 = *; p < 0.01 = **; p < 0.001 = ***.

To confirm that *WNT11^Δ814-1062^* encodes a dominant-negatively acting Wnt ligand, we next injected *Xenopus* embryos from a Wnt/β-catenin signaling reporter line (pbin7LEF::dGFP; (Haas et al., 2019)) unilaterally targeting the right neural plate and analyzed Wnt-driven GFP expression at st. 20 by epifluorescent microscopy. Additionally, we used overexpression of *Xenopus wnt3a* (Kiecker and Niehrs, 2001) as positive control to stimulate reporter activity. In control embryos, reporter activity was bilaterally symmetric, while Wnt3a increased reporter activity as expected (**Fig. S2A,B**). In contrast, overexpression of WNT11^c.814delG^did not profoundly alter reporter activity on the injected side, while WNT11^Δ814-1062^ strongly and WNT11^Δ850-1062^ weakly reduced Wnt reporter activity (**Fig. 4A,B**). These data suggested that removal or replacement of frameshift amino acids from WNT11^c.814delG^ generated dominant-negative acting WNT11 ligands.

**Figure 4:**
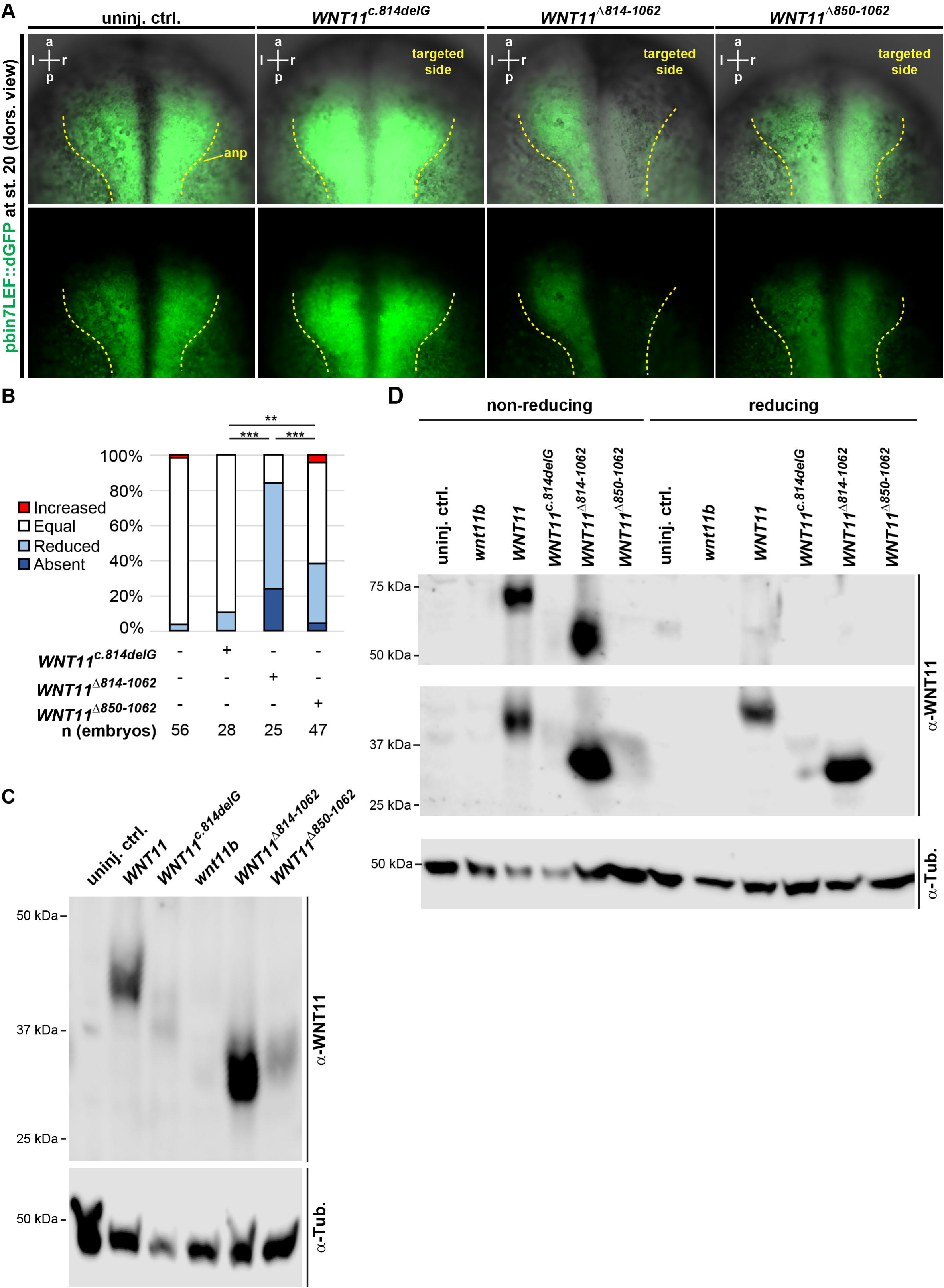
WNT11c^.814delG^ encodes a loss-of-function variant. **(A,B)** Overexpression of mRNAs encoding WNT11^c.814delG^, WNT11^D814-1062^ and WNT11^D850-1062^, and analysis of effects on Wnt/β-catenin signaling reporter (pbinLEF::dGFP) activity (green). **(A)** Representative images. **(B)** Quantification of results. n = number of embryos analyzed per condition. *χ*^2^-test, p > 0.05 = ns; p < 0.05 = *; p < 0.01 = **; p < 0.001 = ***. **(C,D)** Overexpression of mRNAs encoding WNT11, WNT11^c.814delG^, WNT11^D814-1062^, WNT11^D850-1062^ and Wnt11b, and analysis of **(C)** protein expression levels as well as **(D)** dimerization of ligands using Western blotting. α-WNT11 = anti-human WNT11 antibody; α-Tub. = anti-alpha-Tubulin antibody (loading control). Full membranes are shown in supplemental figure S2C and D.

The observed functional differences in C-terminally truncated WNT11 ligands could be due to changed molecular signaling properties of the ligands (e.g. loss of dimerization ability) or could lead to altered expression levels, e.g. by triggering protein degradation (e.g. due to protein misfolding). To investigate which scenario applied to WNT11^c.814delG^, we overexpressed WNT11, WNT11^c.814delG^, WNT11^Δ814-1062^ or WNT11^Δ850-1062^ and assessed protein levels as well as dimerization using sodium dodecyl-sulfate polyacrylamide gel electrophoresis (SDS-PAGE) and Western blotting. To ensure specific reactivity of the anti-WNT11 antibody to the overexpressed human WNT11 constructs, we included uninjected as well as *wnt11b* injected controls in these assays. Uninjected and *wnt11b* injected samples showed only minor reactivity to the anti-WNT11 antibody, while a clear band around the expected 40 kDa was detected in WNT11-injected samples (**Fig. 4C; S2C**). In contrast to WNT11, WNT11^c.814delG^ and WNT11^Δ850-1062^ showed reduced abundance (**Fig. 4C; S2C**), linking their reduced functional potencies to reduced protein concentrations. The modified ligand WNT11^Δ814-1062^, which also showed strong dominant-active activity, was in turn detected at the expected size (around 30 kDa) and at levels similar to the wt WNT11 ligand (**Fig. 4C; S2C**). Comparison of Western results between reducing (standard) vs. non-reducing (allowing to detect Wnt ligand-dimers; (Cha et al., 2009)) sample preparation conditions further indicated that all detectable ligands, including WNT11 and WNT11^Δ814-1062^, were able to form ligand dimers *in vivo* (**Fig. 4D; S2D**), a prerequisite for signaling activity.

Thus, we conclude from these results that changes in the last 12 amino acids in WNT11^c.814delG^ render the ligand instable, leading to reduced protein levels and loss of normal or dominant-negative signaling activity.

## Discussion

Our functional studies to characterize the novel human *WNT11^c.814delG^* variant revealed that it encodes a loss-of-function allele: Overexpression of *WNT11^c.814delG^* did not affect morphogenesis, left-right axis patterning and canonical Wnt signaling activity in *Xenopus* embryos. This was surprising, because the variant encodes a ligand with nearly identical length and predicted structure to the *Xenopus* dominant-negative Wnt11b construct (283 vs. 282 amino acids respectively) as well as to a dominant-negative acting Wnt11b ligand resulting from mis-splicing with 296 amino acid length (Dichmann et al., 2015; Tada and Smith, 2000). This lack of activity is not caused by functional differences of Wnt11 homologs across species, because overexpression of full-length WNT11 yielded similar results to *Xenopus* Wnt11b. Instead, our results suggest that the functional differences are due to changes of the composition of the last 12 frameshift amino acids. Removal of the 12 frameshift amino acids (*WNT11^Δ814-1062^*) restored dominant-negative activity. Moreover, replacing these 12 amino acids with the wt sequence (*WNT11^Δ850-1062^*) resulted in a ligand with weaker, yet clear dominant-negative activity. Our results further indicate that the lack of signaling activity by the WNT11^c.814delG^ variant is due to reduced protein levels, while C-terminally truncated ligands can generally still dimerize (Cha et al., 2008).

Interestingly, removal of 12 amino acids from *Xenopus* dnWnt11b (*Wnt11b^Δ811-1059^*) also resulted in a weaker dominant-negative activity of the ligand, and it was described that dominant-negative activity of Wnt ligands depends on their length (Hoppler et al., 1996; MacDonald et al., 2014). The first dominant-negative Wnts were constructed by C-terminally truncating the ligand behind a particular conserved cysteine (C13). In search of a dominant-negative Wnt8, the authors noted that a 30 amino acids shorter ligand than dnWnt8, which lacked the conserved cysteine, had only weak dominant-negative activity (Hoppler et al., 1996). Subsequently, dnWnt11b was designed with the premature stop positioned directly after the conserved cysteine residue (Tada and Smith, 2000). Nevertheless, our results reveal that dominant-negative behavior of Wnts cannot be dependent on a specific length or the presence of the cysteine residue, as WNT11^Δ814-1062^ and Wnt11b^Δ811-1059^ do not contain it. Instead, we propose that a combination of ligand length and amino acid composition is required for protein stability and dominant-negative activity, perhaps by allowing certain posttranslational modifications, including glycosylation and acylation (Willert and Nusse, 2012), which are indispensable for Wnt structure and secretion (Komekado et al., 2007; Kurayoshi et al., 2007). Amino acid changes might increase susceptibility for recognition by protein control mechanisms, such as unfolded protein response or destabilization (Teng et al., 2010). In contrast, nonsense-mediated decay of the mRNA seems rather unlikely, because we overexpressed synthetic mRNAs at high levels and the stop codon in *WNT11^c.814delG^* is positioned 120 bases upstream of the last exon-exon-junction, a region which is typically insensitive to nonsense-mediated decay (Nagy and Maquat, 1998).

In line with our experimental results, the affected individual is homozygous for *WNT11^c.814delG^*, while both heterozygous parents were healthy individuals (except for SIT in the mother), arguing against a dominant-negative function of WNT11^c.814delG^in humans. The phenotypic features observed in the affected child, including TOF, renal hypodyplasia and SIT match phenotypic features previously identified in *Wnt11^-/-^* mice, e.g. CHDs, including transposition of the great arteries, double outlet right ventricle, persistent truncus arteriosus, and ventricular septum defects with complete penetrance (Zhou et al., 2007). These CHDs all belong to the group of outflow tract defects, which also includes TOF, defined as a combination of pulmonary stenosis, ventricular septal defects (VSD), overriding aorta and hypertrophy of the right ventricle (Neeb et al., 2013). *Wnt11^-/-^* mice also displayed hypoplastic kidneys, where the tips of ureteric buds were partially lost and ureteric branching was affected (Majumdar et al., 2003). Nagy et al. observed renal hypoplasia in 25% of *Wnt11* knockout animals with additionally reported secondary glomerular cysts. Lack of WNT11 resulted in reduced cell proliferation and increased apoptosis in the cortex as well as reduced expression of *Wnt9b*, *Six2, Hox10 and Foxd1* at E16.5. Additionally, they observed reduced tubular convolution, prompting the authors to hypothesize that disturbed conversion extension movements may contribute to the observed renal phenotype (Nagy et al., 2016). *Wnt11b^-/-^ Xenopus* showed laterality defects, in line with a previous study on the role of Wnt11b in *Xenopus* (Houston et al., 2022; Walentek et al., 2013). These results, generated in model organisms in combination with the patient case presented in this study, strongly suggest similar functions of WNT11 in human development.

Furthermore, biallelic loss-of-function of WNT11 should be considered as novel underlying genetic cause in syndromal human phenotypes presenting with congenital heart defects and renal hypoplasia / dysplasia, with or without laterality defects. Congenital anomalies of the kidneys and urinary tract (CAKUT) and cardiac malformations are known to co-occur in a number of human syndromes, many of which result from chromosomal aberrations. However, bilateral renal hypoplasia overall is rarely observed. Interestingly, autosomal dominantly inherited *WNT5a* dysfunction also results in a right ventricular outflow tract obstruction and renal anomalies with incomplete penetrance in addition to skeletal dysplasia features (Robinow syndrome (OMIM#180700) (Person et al., 2010). Wnt5 ligands share functional similarities with Wnt11 ligands, and in mice, *Wnt5a* loss-of-function likewise causes renal developmental defects, including uni- or bilateral kidney agenesis and hypoplasia with altered pattern of ureteric tree organization as well as duplex kidneys with reduced penetrance (Pietilä et al., 2016). It has also been suggested that both WNT11 and WNT5a are required for proper secondary heart field development where they govern non-apoptosis related caspase activities and promote cardiac progenitor development (Bisson et al., 2015).

It remains elusive why the heterozygous mother of the affected boy presented with SIT. Functional haploinsufficiency during laterality development of the mother in combination with additional stressors could have contributed to the development of SIT, although there is no indication for haploinsufficiency in heterozygous *Xenopus wnt11b* mutants (Houston et al., 2022). While whole exome sequencing of the mother did not reveal any putative genetic cause for laterality defects, genetic abnormalities are generally identified in less than half of cases (Antony et al., 2022). Therefore, the mother could coincidentally have a SIT independent of the *WNT11* variant.

In conclusion, this work revealed *WNT11* dysfunction as a novel likely cause of a complex developmental phenotype in humans, and elucidated functional properties of Wnt ligands *in vivo*. This enhances our understanding of the structure-function relationship in Wnt ligands with clinical relevance.

## Acknowledgments

We thank: S. Schefold for expert technical help; S. Arnold for support and discussions; C. Softley for help with setting up Western blots; Xenbase, EXRC for Xenopus resources; Light Imaging Center Freiburg, BiMiC and Aqua Core for microscope/animal resources, and B. Grüning and the Freiburg Galaxy Team for bioinformatics platform and support. This study was supported by the Deutsche Forschungsgemeinschaft (DFG) under the Emmy Noether and Heisenberg Programmes (grant WA3365/2-1 and WA3365/5-1), to PW; and by DFG SFB1453 NephGen (Project ID 431984000) as well as under Germany’s Excellence Strategy (CIBSS - EXC-2189 - Project ID 390939984) to PW and MS. MS also acknowledges funding from the European Research Council (ERC; starting grant TREATCilia grant # 716344) and SFB1597 SmallData (Project ID 503306912). HB was supported by the MOTI-VATE program for medical scientists, Faculty of Medicine, University Freiburg.

## Author contribution

HB: Xenopus experiments; MH, MMBE, PW: Xenopus experimental support; HB, MS, PW: experimental design, planning, analysis and interpretation of data; ZB, MS: Human subject recruitment, clinical classification and human exome sequencing / analyses. ZB: human Sanger sequencing and analyses. HB, PW, MS: manuscript preparation with input from all authors. MS, PW: study design and supervision, coordinating collaborative work.

## Figure legends

**Figure S1: Sequence comparison and mortality rates**

**(A)** Alignment of nucleotide sequences of *WNT11*, *WNT11^c.814delG^*, *Wnt11b* and *dnWnt11b*. **(B)** Mortality rates recorded in experiments corresponding to main figure 1A,B. **(C)** Alignment of amino acid sequences of WNT11, WNT11^c.814delG^, Wnt11b and dnWnt11b.

**Figure S2: Validation of Wnt-reporter sensitivity and Western blot membranes**

**(A)** Overexpression of *wnt3a* mRNA and analysis of effects on Wnt/β-catenin signaling reporter (pbinLEF::dGFP) activity (green). **(A)** Representative images. **(B)** Quantification of results. n = number of embryos analyzed per condition. **(C)** Full membranes of experiments corresponding to main figure 4C. **(D)** Full membranes of experiments corresponding to main figure 4D.

## Material and Methods

### Identification of patient variant using exome sequencing

DNA samples were collected after obtaining informed consent from affected individuals or parents as part of the clinical diagnostic pathway at Radboud UMC Nijmegen (Innovative diagnostic program). Human subjects research was approved by the Ethics committee of the Institute of Child Health, University College London, London UK (GOSH R&D number 11MM03, REC number 08/H0713/82) the ethics committee Arnhem-Nijmegen, The Netherland (ethical approval no 2006–048) and the ethics committee of Freiburg University, Freiburg, Germany (votum no 122/20).

Exome sequencing and sequence analysis was performed as described previously (Loges et al., 2018). Exomic sequences from DNA samples were enriched using SureSelect Human All Exon V.6 Kit (Agilent Technologies #5190-8892) according to the manufacturerer protocol followed by sequencing on a Hiseq PE150 (Illumina, San Diego, California, USA). Read alignment and variant calling were performed using Burrows-Wheeler (BWA)/GATKpipeline using default parameters with the human genome assembly hg19 (GRCh37) as reference. Vtools and ANNOVAR softwares were used to store and annotate variants. Following alignment and variant calling, serial variant filtering was performed for variants with a MAF equal or less than 1% in ExAc, 1000 genome project and esp6500 and gnomAD databases, coding variants or variants within 5 bp of exon-intron boundaries. Obligate loss-of-function variants such as canonical splice variants, frameshift and stop mutations were prioritized over missense variants, however missense variants were not excluded from the analysis. CNV calling was performed using Exome Depth (Plagnol et al., 2012) and BAM files were visually inspected for homozygous CNVs in all genes known to cause laterality defects or renal hypo(dysplasia). All the variants passing these filters were subsequently inspected at aligned read level with the aim of avoiding false call due to misalignment or low-depth of coverage.

### Sanger Sequencing

Genomic DNA was isolated by standard methods using a Qiagen kit for blood samples. Genomic DNA amplification was performed in a volume of 50 µl containing 30 ng DNA, 50 pM of each primer, 2 mM dNTPs, and 1.0 U GoTaq DNA polymerase (Promega, #M3001). PCR amplifications were carried out by an initial denaturation step at 94°C for 3 min, and 33 cycles as follows: 94°C for 30 sec, 58-60°C for 30 sec, and 72°C for 70 sec, with a final extension at 72°C for 10 min. PCR products were verified by agarose gel electrophoresis, purified, and sequenced bi-directionally. Sequence data were analysed using the CodonCode software. Primer sequences are listed below.

Primers used for sanger sequencing (5’-3’):

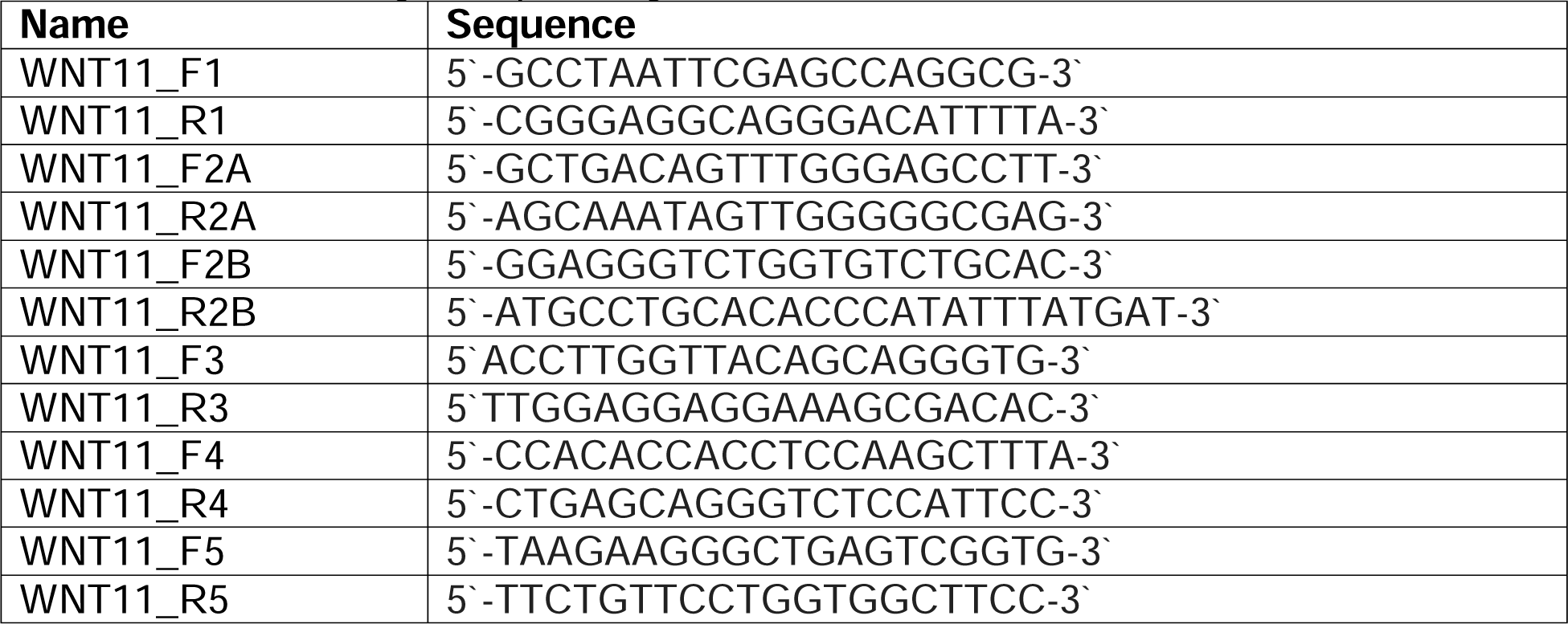

### Human data availability

The datasets for this article are not publicly available due to concerns regarding participant/patient anonymity. Requests to access the datasets (e.g. a pre-filtered variant list) should be directed to.

### Protein structure prediction

Protein structure predictions were created on the Galaxy EU platform (https://usegalaxy.eu/), using Alphafold 2 (v.2.1.2) (toolshed.g2.bx.psu.edu/repos/galaxy-australia/alphafold2/alphafold/2.1.2+galaxy4) (Community, 2024; Jumper et al., 2021) and adding the amino acid sequences into the Fasta Input. For each protein, “Model 1” from the top five predicted structures was selected. Models were displayed in Cartoon or Surface representations and snapshots of the models were taken from approximately the same angles.

### MultAlin amino acid alignments

Alignments of amino acid sequences were made using MultAlin (multalin.toulouse.inra.fr/multalin (Corpet, 1988)) and the following symbol comparison Table: Blosum62-12-2. High consensus value (red) was set to 90% and low consensus value (blue) to 50%.

### Animal experiments

Wild-type and transgenic *Xenopus laevis* were obtained from the European Xenopus Resource Centre (EXRC) at University of Portsmouth, School of Biological Sciences, UK, or Xenopus 1, USA. Frog maintenance and care was conducted according to standard procedures in the AquaCore facility, University Freiburg, Medical Center (RI_00544) and based on recommendations provided by the international Xenopus community resource centers NXR (RRID:SCR_013731) and EXRC as well as by Xenbase (http://www.xenbase.org/, RRID:SCR_003280)(Fisher et al., 2023). This work was done in compliance with German animal protection laws and was approved under Registrier-Nr. G-22/43 by the state of Baden-Württemberg.

### Animal experimental data availability

Imaging and quantification data are available to the scientific community upon request to.

### Manipulation of *Xenopus* Embryos

*X. laevis* eggs were collected and in vitro-fertilized, then cultured and microinjected by standard procedures (Sive et al., 2007; Sive et al., 2010). Embryos were injected with mRNAs at four-cell to eight-cell stage using a PicoSpritzer setup in 1/3x Modified Frog Ringer’s solution (MR) with 2.5% Ficoll PM 400 (GE Healthcare, #17-0300-50), and were transferred after injection into 1/3x MR containing Gentamycin. Drop size was calibrated to about 7–8nL per injection.

Human *WNT11* (*HWNT11-pCMV6-Entry*) was obtained from OriGene (#RC219688) (matching ENST00000322563.8, NM_004626.3, CCDS8242) and subcloned into pCS108 (using Cla1 and EcoR1 enzymes; NEB, #R0197 and #R3101) to serve as templates for *in vitro* mRNA synthesis using primers listed below. *Xenopus* Wnt11b and dnWnt11b constructs were derived from (Dichmann et al., 2015; Tada and Smith, 2000). *Xenopus* Wnt3a encoding construct was a gift from C. Niehrs (Kiecker and Niehrs, 2001). All variants were generated by site-directed mutagenesis (NEB, #E0554S) using primers listed below. mRNAs encoding WNT11, WNT11c.814delG, WNT11^Δ814-1062^, WNT11^Δ850-1062^, Wnt11b, dnWnt11b and Wnt11b^Δ811-1059^ (70-100 ng/μl), Wnt3a (5 ng/μL) were injected together with *centrin4-gfp* (50 ng/μl) or Rhodamine Dextran (Invitrogen, #11590226) as lineage tracers. All mRNAs were prepared using the mMessage Machine kit using Sp6 (Invitrogen, #AM1340) supplemented with RNAse Inhibitor (Promega, #N251B).

Cloning primers to generate new WNT11 and Wnt11b constructs (5’-3’):

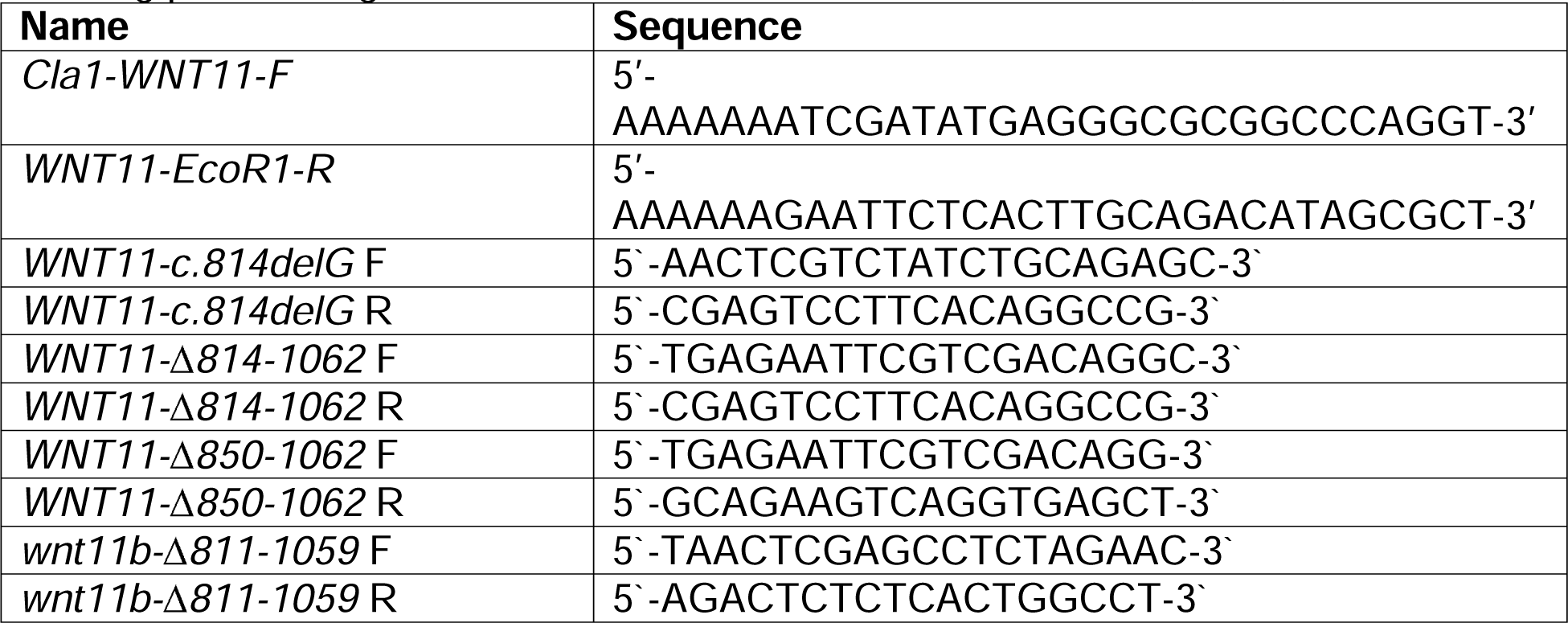

### Whole mount *in situ* hybridization

For whole mount *in situ* hybridization against *pitx2c,* the antisense probe was derived from (Schweickert et al., 2001). Embryos were fixed in MEMFA (100mM MOPS pH7.4, 2mM EGTA, 1mM MgSO4, 3.7% (v/v) Formaldehyde) overnight at 4°C and stored in 100% Ethanol at -20°C until used. DNAs were purified using the PureYield Midiprep kit (Promega, #A2492) and were linearized before in vitro synthesis of anti-sense RNA probes using T7 or Sp6 polymerase (Promega, #P2077 and #P108G), RNAse inhibitor and dig-labeled rNTPs (Roche, #3359247910 and 11277057001). Embryos were *in situ* hybridized according to (Harland, 1991), and stained with BM Purple (Roche, #11442074001) and imaged.

### Evaluation of morphology, WMISH staining, and Wnt reporter activity

Embryos were staged according to Nieuwkoop and Faber (1994) Normal Table of *Xenopus laevis* (Daudin) (Zahn et al., 2022) and Garland Publishing Inc, New York ISBN 0-8153-1896-0. Embryos that could not be assessed for *pitx2c* expression patterns (due to missing axes or coiled morphology) were not included in the statistics for *pitx2c* expression. Images of embryos for morphological evaluations and after *in situ* hybridization were generated using Zeiss Stemi508 with Axiocam208-color, and images were adjusted for color balance, brightness and contrast using Adobe Photoshop.

Canonical Wnt-reporter activity was assessed in transgenic embryos (pbin7Lef::dGFP; line: Xla.Tg(WntREs:dEGFP)^Vlemx^)(Haas et al., 2019) by comparing fluorescence between the manipulated and non-manipulated control side on imaged generated using a Zeiss AxioZoom setup. Images were adjusted for brightness and contrast using imageJ (Schindelin et al., 2012) and Adobe Photoshop.

### SDS PAGE and Western blotting

8-15 embryos were placed in an Eppendorf tube without fluid and stored at -20 °C. For the standard Western Blot, 100 µl of 1x Lysis Buffer were added (20 mM Tris-HCl pH8, 150 mM NaCl, 2 mM EDTA, 1x Protease Inhibitor Roche, #04693116001, 1% NP40 Sigma, #I8896), the embryos were smashed by pipetting up and down several times and the samples were centrifuged at 4 °C at maximum speed for 15 minutes. For the dimerization assay, as has been described by (Cha et al., 2009), embryos were instead smashed by pipetting up and down with 90 µl ice-cold PBSt and 10 µl Protease Inhibitor (Roche, #04693116001) and the samples were centrifuged at 4 °C at 14400 g for 10 minutes.

The supernatant was transferred to a fresh Eppendorf tube. 4x Laemmli Buffer (50 ml 4x buffer, 1 M Tris, pH 6.8, 4 g SDS, 20ml Glycerol, 10 ml 2-Mercaptoethanol, 0.1 g Bromophenol Blue) was added to the supernatant to have a 1x final concentration. For the standard Western Blot, 4x Laemmli Buffer containing 5 % 2-Mercaptoethanol (Roth, #4227.2) was used and the samples were cooked at 95 °C for 5-10 minutes on a shaking plate. For the Dimerization Assay, 4x Laemmli Buffer without 2-Mercaptoethanol (non-reducing condition) or 10 % 2-Mercaptoethanol (reducing condition) was used, and the samples were not cooked.

For 2 gels, 10 ml of 8 % Separating Gel was made of: 2.5 ml 4x Tris SDS (Roth, #2326), pH 8.8, 2 ml 40 % Acrylamide (Sigma, #A7802), 5.4 ml H_2_O, 40 µl TEMED (Roth, #2367.1), 100 µl 10 % APS (Roth, #9592.2). For 2 gels, 5 ml of 4 % Collecting Gel was made of: 1.25 ml 4x Tris SDS, pH 6.8, 0.625 ml 40 % Acrylamide, 3.11 ml H_2_O, 50 µl TEMED, 50 µl 10 % APS). 1x Running buffer (25 mM Tris-HCl, pH 8, 192 mM Glycine (Roth, #3187) in distilled water) was used for electrophoresis. 10-12 µl Precision Plus Protein Western C Standards Ladder (BioRad; #161-0376) and 10-20 µl of each sample were loaded. The electrophoresis was run at 120 V for 1-2 hours at RT.

Semi-dry transfer onto an activated PVDF membrane (Thermo Scientific, #88518) was conducted in 1x Towbin buffer with 0.1 % SDS for 30 minutes (25 mM Tris-base, 192 mM Glycine, 1 % SDS) using a PerfectBlue Semi-Dry Electroblotter Sedec M (VWR; 700-1220). Transfer was performed at a constant current of 10 mA for 90 minutes. Membranes and Gels were stored in 1x TBStw (100 mM Tris-base, 500 mM NaCl, 1 % Tween 20) at 4 °C until further use.

Membranes were blocked for at least 45 minutes using 5 % non-fat dry milk (Roth, #T145.3). The following primary antibodies were used at 1:1000 and incubated over night at 4 °C: Rabbit Polyclonal Anti-WNT11 (ThermoFisher, #PA5-21712) and, as a loading control, Mouse Monoclonal Alpha-Tubulin (Cell Signalling, #3873). The membrane was then washed in 1x TBStw for 4x 20 minutes. The following secondary antibodies were used at 1:3000 and incubated for 2 h at RT: HRP-linked Anit-Mouse IgG (Cell Signalling, #7076) and HRP-linked Anti-Rabbit IgG (Cell Signalling, #7074). Afterwards, the membrane was washed in 1x TBStw for 6x 10 minutes. Membranes were incubated with a mixture of 500 µl of Peroxide solution and 500 µl of Luminol/enhancer solution (both from Clarity™ Western ECL Substrate (BioRad, #170-5061) for 5 minutes in the dark at room temperature. Membranes were imaged using the Odyssey XF Imaging System by LI-COR. Afterwards, membranes were washed and stored in TBStw. Membranes were stripped for loading control (α-Tubulin) re-probing with Acid Stripping Buffer (for 100 ml: 5 ml 2 M Glycine-HCl, pH 2.1, 500 µl Tween 20, 700 µl 2-Mercaptoethanol) for 1 h on a tumbler. The membranes were again rinsed in distilled water and washed 3x 5 minutes with TBStw. To validate effective membrane stripping, the membranes were imaged before re-probing.

### Statistical evaluation

Stacked bar graphs were generated in Microsoft Excel. Sample sizes for all experiments were chosen based on previous experience and used embryos derived from at least two different females. No randomization or blinding was applied. For statistical evaluation, chi-squared (χ^2^) test was performed using Microsoft Excel.

### Use of shared controls

For some of the *in situ* and morphological analyses the same embryos were used in multiple graphs.

