## Supplementary figures and images for "A homozygous human *WNT11* loss-of-function variant associated with laterality, heart and renal defects"

### Fig. S1

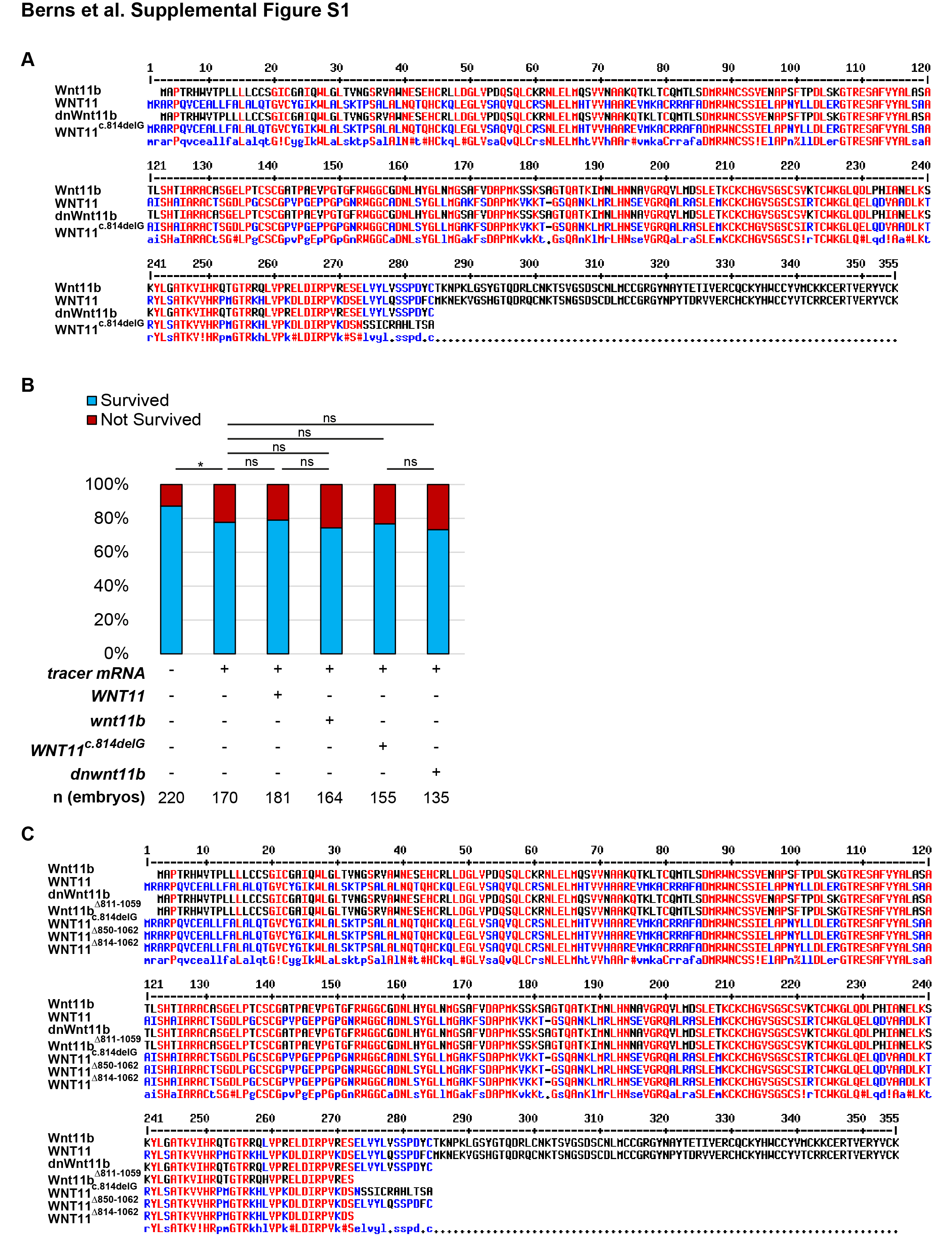

### Fig. S2

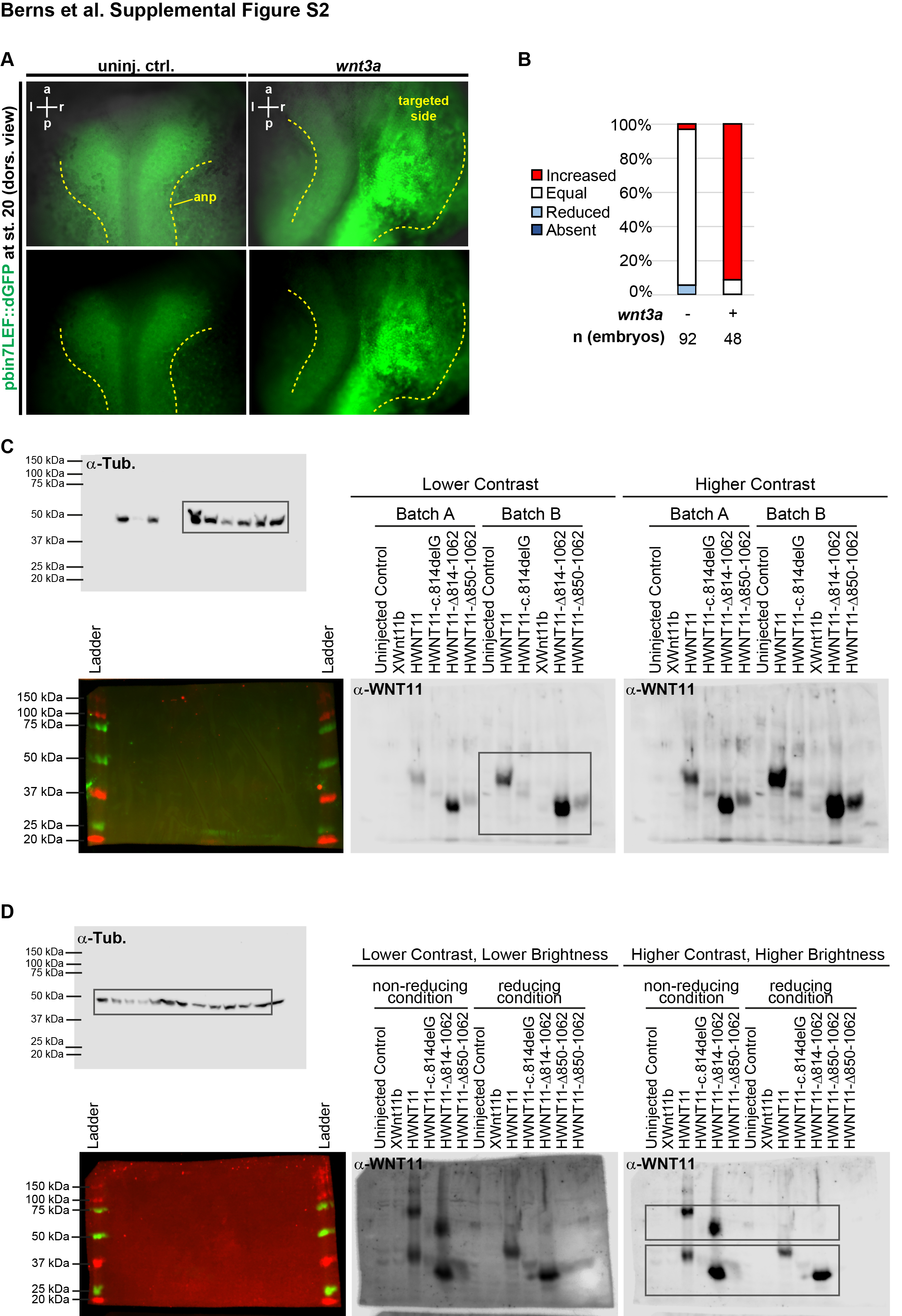
